# Efficient and robust NK-Cell transduction with Baboon Envelope pseudotyped lentivector: a major tool for immunotherapy

**DOI:** 10.1101/625285

**Authors:** Aurelien B.L. Colamartino, William Lemieux, Panojot Bifsha, Simon Nicoletti, Nitin Chakravarti, Joaquin Sanz Remon, Hugo Roméro, Silvia Selleri, Kathie Béland, Mélanie Guiot, Camille Tremblay-Laganière, Renée Dicaire, Luis Barreiro, Dean A. Lee, Els Verhoeyen, Elie Haddad

## Abstract

NK-cell resistance to transduction is a major technical hurdle for developing NK-cell immunotherapy. By using Baboon envelope pseudotyped lentiviral vectors (BaEV-LVs) encoding eGFP, we obtained a transduction rate of 23.0±6.6% in freshly-isolated NK-cells (FI-NK) and 83.4±10.1% in NK-cells obtained from the NK-cell Activation and Expansion System (NKAES), even at low MOI, with a sustained transgene expression for at least 21 days. BaEV-LVs outperformed Vesicular Stomatitis Virus type-G (VSV-G)-, RD114-and Measles Virus (MV)-pseudotyped LVs (p<0.001). mRNA expression of both BaEV receptors, ASCT1 and ASCT2, was detected in FI-NK and NKAES, with much higher expression in NKAES. Transduction with BaEV-LVs encoding for CAR-CD22 resulted in robust CAR-expression on 44.2%±14.2% of NKAES cells, which allowed the specific killing of the NK-resistant pre-B-ALL-RS4;11 cell line. Using a larger vector, encoding a dual CD19/CD22-CAR separated by T2A, we were able to transduce and re-expand dual-CAR-expressing NKAES, even with low viral titer. These dual-CAR-NK efficiently and specifically killed both CD19^KO^-and CD22^KO^-RS4;11 cells, which may overcome antigen-loss escape in the clinical setting. Our results suggest that BaEV-LVs may efficiently enable NK-cell biological studies and translation of NK-cell-based immunotherapy to the clinic.

## INTRODUCTION

The relative resistance of NK cells to transduction hampers the study of NK-cell biology and the development of NK cell-based immunotherapy. VSV-G-LVs, classically used to generate chimeric antigen receptor (CAR)-T cells,(1) do not efficiently transduce NK-cells. RD114-pseudotype viral vectors represent an attractive alternative since their entry receptor, the sodium-dependent neutral amino acid transporters (ASCT2),(2, 3) is widely expressed in the hematopoietic lineage.(4) However, despite encouraging initial report,(5) RD114-based viral vectors only transduce NK-cells at low levels. Nevertheless, they recently enabled clinical development of cord-blood derived CAR-NK-cells.(6) To fill the need for an efficient method for transducing NK cells, we investigated alternative pseudotyping proteins. Since the Baboon envelope pseudotyped lentiviral vector (BaEV-LV) binds ASCT1(7) in addition to ASCT2 for viral entry, we assessed their efficacy to transduce NK-cells for therapeutic purposes.

## RESULTS AND DISCUSSION

We first transduced NK cells expanded using the Amplification and Expansion System (NKAES) and freshly isolated NK-cells (FI-NK) with an eGFP-encoding LV and observed that in both cases, BaEV-LVs outperformed VSV-G-, MV-, and RD114-LVs (Figure 1A, 83.4 % mean transduction rate vs 15.7%, 13.7% and 37.8% for NKAES, p <0.0001, and 23.0% vs 10.4%, 2.1% and 7.8% for FI-NK, p <0.0001, respectively). The mean transduction rate with BaEV-LVs was higher than 60% for NKAES even at low MOI of 1, and ranged from 12.4% at a MOI of 1 to a maximum of 27.2% at a MOI of 10 for FI-NK (Figure 1B). The number of living cells in both NKAES and FI-NK was preserved after transduction with BaEV-LVs (Figure 1C), suggesting that transduction did not affect viability or NK-cell proliferation. Transgene expression persisted over time after transduction with BaEV-LVs, although a decrease was observed from 70.6% to 61.4% in 14 days for NKAES (p=0.06). Transduced FI-NK could be easily amplified after transduction (not shown). High transduction rates were also observed after NK-cell expansion on K562-mbIL15-41BBL feeder cells(9) or feeder-free NK MACS medium(15) (Figure 1D).

**Figure 1.**
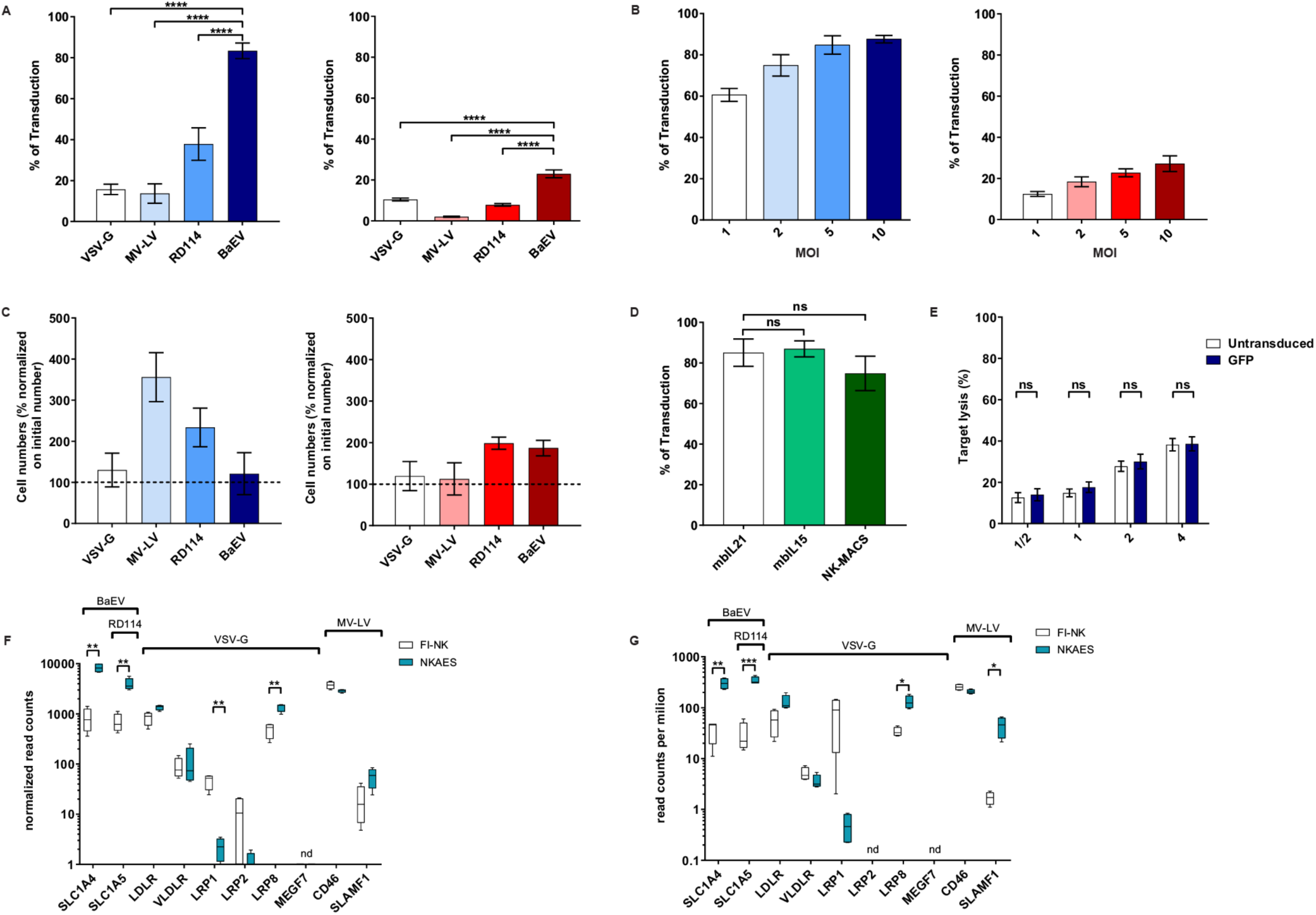
BaEV-LV efficiently transduces expanded (NKAES) and freshly isolated NK-cells (FI-NK). **(A)** Transduction of NKAES cells (K562mbIL15 system, left panel) or FI-NK (right panel) using VSV-G (n=3 and n=6), Measles virus (MV; n=3), RD114 (n=7 and n=6) or BaEV (n=7 and n=12) envelope pseudotyped LVs encoding eGFP. Fluorescence was evaluated by flow cytometry (P<0.001; one-way ANOVA test with Bonferroni correction). **(B)** Transduction of NKAES (left panel; n=5) or FI-NK (right panel; n=3) using BaEV-LV-encoding eGFP at various multiplicities of infection (MOI) was measured by fluorescence.**(C)** Viability of NKAES (left panel; n=3) or FI-NK (right panel; VSV-G and MV-LV n=6, RD114 n=3, BaEV n=9) with different LVs. Percentage of living cells in the culture are normalized to the initial number of cells before transduction. **(D)** BaEV-LV transduction with NK Amplify and Expansion System (NKAES) using K562mbIL21 or K562mbIL15 irradiated feeder cells (n=4), or using the NK-MACS Medium (P= n.s.; one-way ANOVA test with Bonferroni correction, n=3). **(E)** Cytolytic function of BaEV-LV-transduced vs untransduced NKAES cells against K562 target cells. Data represent the percentage of target lysis, assessed by flow cytometry relative to the effector/target ratio and normalized to targets alone. (n=3; p=n.s.; 2-way ANOVA test with Bonferroni correction). (**F**) Quantitative expression of the viral receptors mRNA assessed by RNA-seq in NKAES expanded with K562mbIL21 and in FI-NK-cells (n=4; *p<0.05; **p<0.01, ***p<0.001, T test with Holm-Sidak correction). (**G**) Quantitative expression of the viral receptors mRNA assessed by RNA-seq in NKAES expanded with K562mbIL15 and in FI-NK-cells (n=4; *p<0.05; **p<0.01, ***p<0.001, T test with Holm-Sidak correction). Data are presented as the mean ± SEM.. For all experiments, NKAES and FI-NK were assessed at day 3 or day 5 post-transduction, respectively.

We then assessed whether NK-cell cytotoxic function was preserved after BaEV-LV transduction and confirmed that the cytotoxicity of eGFP-transduced NKAES cells against K562 cells was equivalent to non-transduced NKAES (Figure 1E).

RNAseq analyses of both FI-NK and NKAES showed that ASCT1 and ASCT2 mRNAs were detected at significantly higher frequency in both IL-15-and IL-21-NKAES than in FI-NK (Figure 1F-G), which may explain the higher transduction rate of NKAES. These data were confirmed by qPCR (not shown). Also, the expression of both BaEV receptors by NK cells may explain the higher transduction efficacy of BaEV-LVs as compared to RD114 wich use only one of those receptors.

CAR-expressing NK-cells represent one of the most relevant clinical applications of efficient NK-cell transduction. We first tested a single 3^rd^ generation CAR construct recognizing CD22 (Figure 2A). We obtained a transduction rate of 44.2%±14.2% of NKAES sustaining high level of CAR-expression (Figure 2B-C), and we demonstrated that CD22-CAR-NK-cells efficiently and specifically killed B-ALL RS4;11 target cells, which were resistant to untransduced NKAES (Figure 2D). We could obtain 5×10^8^ CAR-expressing cells from 5×10^5^ transduced cells after an expansion of 14 days (not shown). Since transgene size affects transduction efficacy,(16, 17) we also tested a dual CAR-expressing vector with two independent chains recognizing CD19 and CD22. The length of the dual CAR-CD22/19 LV had a significant impact on virus production since titers were under the detection limit (not shown) and NK transduction was very low (<1%). We sorted and re-expanded these dual-CAR-transduced NK-cells for two weeks, to assess their cytotoxicity. Importantly, we showed that these CAR-CD22/19-NKAES killed efficiently CD19^KO^-CD22^KO^-RS4;11 cells, which suggest that this strategy could be efficient for preventing tumor evasion to CAR therapy(18, 19) (Figure 2E).

**Figure 2.**
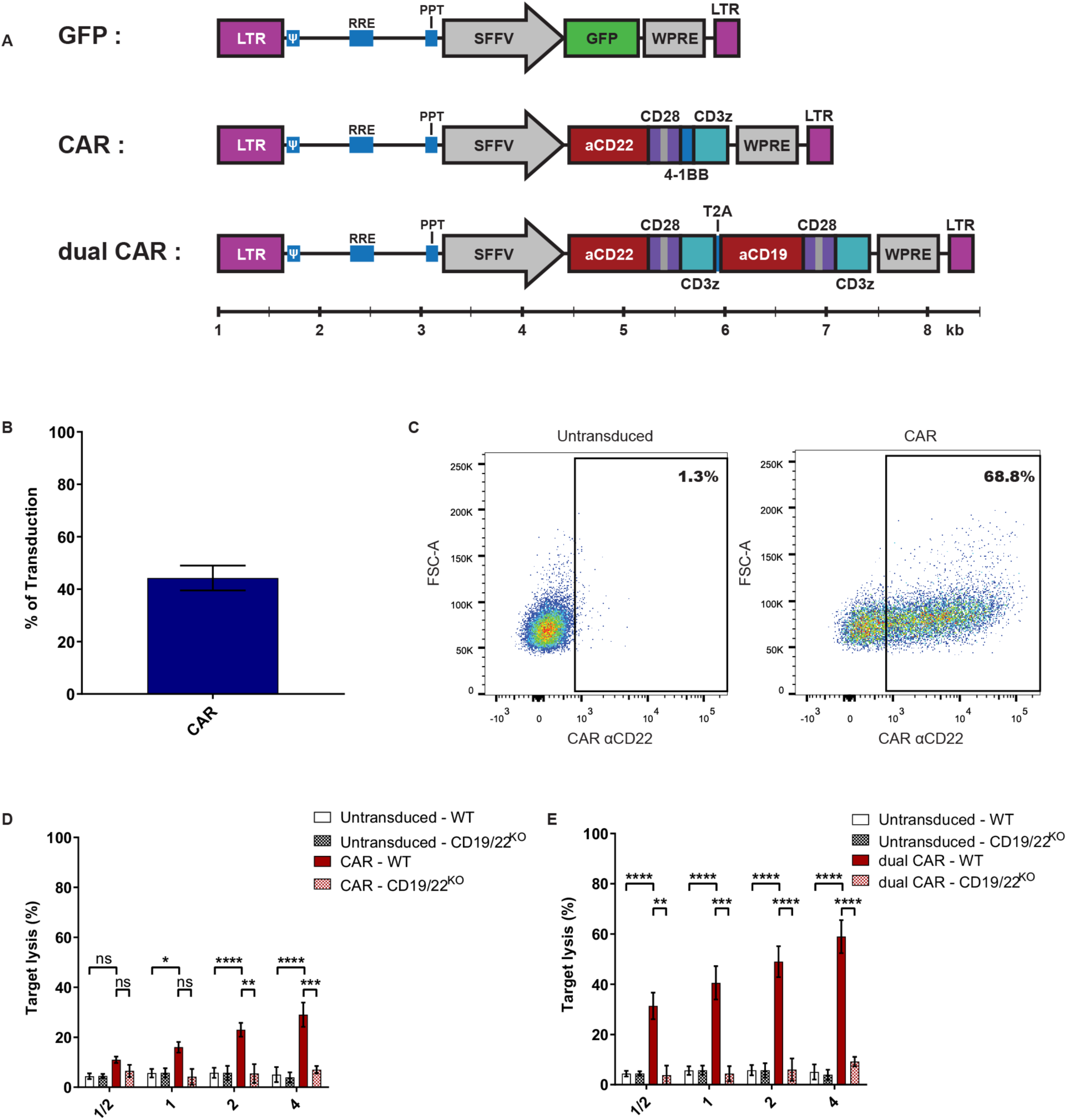
BaEV-LVs allow robust CAR-expression in NK-cells. **(A)** Schematic representation of the different LVs used for NK-cell transduction using BaEV-LV (relative scale according to size in base pairs). **(B)** Percentage of transduced NKAES cells using BaEV-LVs coding for an anti-CD22 CAR (n=9). **(C)** Flow cytometry plot representative of CAR-CD22 expression after NK-cell transduction with BaEV-LVs. **(D)** Cytotoxic assays against either parental (n=7) or CD19/22^KO^-RS4;11 cells (B-ALL; n=2) using CAR-CD22-NK-cells (p<0.0001; 2-way ANOVA test with Bonferroni correction). **(E)** Cytotoxic assays against parental (n=7) or CD19/CD22^KO^ RS4;11 B-ALL cells (n=2) using NK-cells transduced with a dual CAR (3^rd^ construct illustrated in A) (p<0.0001; 2-way ANOVA test with Bonferroni correction). Data are presented as the mean ± SEM. *p<0.05; **p<0.001, ***p<0.0001

In this study we showed that BaEV-LV is an efficient and robust tool to transduce NK cells. As a proof-of-concept, we generated large numbers of engineered CAR-NK-cells, which induced specific killing of antigen-bearing cancer cells, even with a large dual CAR-LV construct. This technique was robust and reproducible in different expansion systems, including a feeder-cell-free system. The higher level of transduction could open up possibilities for the use of this method to generate an immunotherapeutic product. The prevalence of receptors, as seen by RNAseq, could explain the difference seen between the transduction of activated and resting NK cells. The fact that activated NK cells express both entry receptors at high level could also explain why this envelope protein is more efficient than the others. The development of such a tool could have a major impact on both basic research of NK-cell biology study and NK-cell-based immunotherapy.

## METHODS

### Cells and Culture Condition

Blood samples were obtained from healthy volunteers after informed consent (IRB-approved protocol #CER-3527). NK-cells were enriched from PBMC using a CD56-positive selection kit (Stemcell Technologies, Canada). NK cells were expanded using the Amplification and Expansion System (NKAES) with irradiated K562mbIL21 or K562mbIL15 feeder cells as described.(8, 9) Alternatively, NK-cells were amplified using NK-MACS Medium (130-114-429, Miltenyi) system according to manufacturer’s instructions. RS4;11 (ATCC) CD19/22^KO^ were generated using purified Cas9 protein and two gRNA targeting CD19 or CD22 (IDTDNA). CD19^KO^ and/or CD22^KO^ cells were FACS-sorted based on loss of surface marker expression. Cells were cultured in DMEM (Wisent) or RPMI1640 supplemented with 10% FCS and penicillin/streptomycin (Gibco). Media were supplemented with 200 UI/mL IL-2 (Proleukin – Novartis Pharmaceuticals, Canada) for NK-cell cultures.

### Plasmids and Viral production

An UCOE sequence(10) was added to the lentiviral vector pHRSIN-SFFV-eGFP(11) upstream of the SFFV promoter to produce pHUS-GFP vector. For the CAR-expression vector, GFP in pHUS-GFP was replaced by an anti-CD22 CAR (m971 ScFv) fused to 28z and BBz (Dr. Orentas, National Cancer Institute).(12) For the dual CAR-expression vector, the GFP-cassette was replaced by two second-generation anti-CD19 and anti-CD22(10) CARs, separated by a self-cleaving T2A peptide (Figure 2A). The pMD2.G (VSV-G) was a gift from Didier Trono (Addgene plasmid#12259; http://n2t.net/addgene:12259; RRID:Addgene_12259) and pLTR-RD114A(13) (RD114) was a gift from Jakob Reiser (Addgene plasmid#17576; http://n2t.net/addgene:17576; RRID:Addgene_17576). The Measles virus (MV-LV) and BaEV envelope plasmids were used as previously described(7). Titration was performed on HEK293T cells (ATCC) using serial virus dilutions.(7)

### Viral transduction

NKAES were transduced after one week of expansion. A day before transduction, a 12-well plate was coated with RetroNectin (Takara). The following day, concentrated vectors at indicated multiplicity of infection (MOI), were added to coated plates for 4 hours at 37°C. Then NK cells were seeded in these wells in IL-2-supplemented medium and protamine sulfate (Pharmaceutical Partners of Canada Inc) (8 ug/mL). The plates were then centrifuged at 1000g for 1 hour and incubated at 37°C overnight. The next day, IL-2-supplemented medium was added to each well. Transduction was assessed on day 3 or day 5 after transduction for NKAES and freshly isolated NK-cells (FI-NK), respectively.

### Flow cytometry

GFP was detected by flow cytometry on CD56(-APC)^+^ CD3(-PE)^−^ cells (Biolegend). To detect CAR-expression, cells were incubated with 2 µl Siglec2(CD22)-Fc chimera (50 mg/ml, R&D) for 30 minutes at 4°C, washed and stained with anti-Fc-PE (Jackson Immune), anti-CD56-APC and anti-CD3-FITC (Biolegend).

### mRNA quantification

RNA-seq expression studies were independently performed in two laboratories (Accession #GSE128696, #GSE129044). For the FI-NK versus IL-21-NKAES/IL-15-NKAES comparisons, extraction of total RNA was done using the RNeasy mini kit (Qiagen) and Total RNA Purification Plus Kit (Norgen Biotek), respectively. The quality of RNA was verified with 2100 Bioanalyzer (Agilent) prior to preparation of sequencing libraries with the TruSeq RNA Sample Prep v2 Kit. Quality of libraries was verified via Agilent 4200 Tapestation using a High Sensitivity D1000 ScreenTape Assay kit. For the IL-15 NKAES analysis, approximately 60-80 million paired-end 150bp sequence reads per library were generated, whereas for the IL-21 NKAES analysis, 30 million single-end 101 bp sequence reads per library were generated, both using Illumina HiSeq4000 platform. Kallisto, an RNA quantification program based on pseudoalignment was used to obtain read count estimates per gene.(14) The differential gene expression analysis was done using DESeq2, edgeR and limma R packages.

### Statistical analyses

Statistical analyses were performed using GraphPad PRISM 8.0 (GraphPad Software). Statistical significance was determined by one-way or 2-way ANOVA with Bonferoni correction or using simple T tests with Holm-Sidak correction.

## AUTHOR CONTIBUTIONS

A.C., W.L., P.B., S.N., H.R., S.S, and M.G. performed the experiments. A.C., W.L., P.B. and K.B. wrote the manuscript. N.C. and D.A.L. generated the RNAseq data on IL-21 expanded NKAES cells and participated in the redaction of the manuscript. R.D. recruited participants and collected samples. E.H. generated the hypotheses, conceptualized the study and wrote the manuscript. E.V provided BaEVTRless encoding plasmid, discussed results and wrote the manuscript. All authors reviewed and approved the manuscript.

## ACKNOWLEDGEMENTS

The authors would like to thank Dr. Dario Campana (St. Jude Children’s Research Hospital) for the K562-mbIL15-41BBL cell line, and Dr. Rimas J. Orentas from the National Cancer Institute for the CAR-CD22 construct. This work was supported by grants from the Canadian Institutes of Health Research (CIHR) (#PJT-156347 and # 130246), and from the “Fonds d’innovation thérapeutique (FIT)” from the “Fondation Charles Bruneau” to E.H. W.L. is supported by a “Fonds de Recherche en Santé du Québec” (FRQS) scholarship award and A.C. by the Cole Foundation scholarship award. S.N. was supported by the Fondation Bettencourt-Schueller (“Ecole de l’INSERM-Liliane Bettencourt”) and a fellowship from the « Fondation ARC pour la recherche sur le cancer ». E.V. was supported by Lentigen/Miltenyi Inc. (Boro Dropulic) for this study.

